# Revisiting the SRX paradox for myosin and myofibrils

**DOI:** 10.64898/2026.10.27.762302

**Authors:** Kerry Y. Kao, Kalen Z. Robeson, Anthony M. Asencio, Tristan W. Wasley, Timothy S. McMillen, Joseph D. Powers, Michael A. Geeves, Michael Regnier

## Abstract

We recently reported that we could find no evidence of the biochemically-defined super relaxed (SRX) state in tissue-isolated HMM, as defined in the original mant-ATP displacement assay. This is consistent with theoretical predictions that any SRX state would have to be in thermal equilibrium with the disordered-relaxed (DRX) form of myosin heads. We repeat the assay here for isolated myosin, from different sources, in its monomeric form or in synthetic myosin filaments. No slow component of the displacement assay was observed, and we again conclude that there is no evidence for the SRX state for isolated myosins. This does not mean that there are no myosins in the ATPase-inhibiting Interacting Heads Motif (IHM) in our assays, just that any IHM is in rapid equilibrium with DRX heads. Mant-ATP displacement assays in the presence of the inhibitor Mavacamten or the activator 2’-deoxyATP (dATP) do provide evidence of an interaction between the two heads of myosin. In contrast, assays with myofibrils do provide evidence of two components in the mant-ATP displacement assay. We propose that these two components result from distinct populations of myosin within the sarcomere thick filament with different fractions of myosin in the IHM structural state.

## 2. Introduction

We recently reported on multiple problems associated with the mant-ATP displacement assay.^1^ This assay has been widely used to assess the fraction of HMM present in solution as the switched off form, with the heads folded back to interact and inhibit each other’s ability to release the ATP hydrolysis products Pi and ADP (named as the IHM or SRX forms^1–3^). We reported a series of paradoxes resulting from these assays and went on to show that, in contrast with most published reports, for a variety of different tissue-generated HMMs we see only a single-phase displacement reaction. This is predicted to be observed for HMM in which the two forms (SRX and DRX) are in relatively rapid equilibrium, with SRX having a turnover < 10% of that of DRX. The essential conclusion of our work, a single-exponential displacement of mant-ATP interpreted as a rapid equilibrium between SRX and DRX, has also been reported for an expressed form of HMM developed by Don Winkleman with S2 tails of variable length.^4,5^ In our studies, we used HMM prepared from cardiac and skeletal muscle.^1^

In contrast to the above studies with HMM, most (but not all) studies that report two phases two phases in the mantATP displacement reaction use single muscle fibers.^2,6^ It is possible therefore, that the two phases are an emergent property that requires a more complex architecture of the myosin when assembled into filaments in the sarcomere. Here we explore the role of increasing structural complexity, using full-length myosin, either as soluble monomers at 0.5 M KCl or as synthetic thick filaments at 0.1 M KCl as the system closest to the physiological system found in cells. Synthetic thick filaments are small bipolar filaments which retain some features of the muscle thick filament but lack all the non-myosin proteins, significantly titin and MyBP-C. We also compare these with purified myofibrils, which include these additional thick filament proteins.

We report that myosins from a variety of striated muscle types (fast skeletal, slow skeletal or cardiac) all show a single-exponential displacement of mant-ATP, consistent with our previous work on HMM. This is true for monomeric myosins and when myosin is assembled into synthetic filaments. We also report that the concentration dependence of myosin and HMM inhibition by Mavacamten and activation by dATP both suggest cooperativity in the two heads not seen in S1. Finally, and importantly, we find that a two phase mant-ATP displacement is present in myofibrils isolated from porcine ventricular muscle, in agreement with our previous report.^7^ Combined the data suggest that sufficient structural features, not found in most reductionist preparations or constructs, are required for myosin to exist in two or more biochemical and/or structural states.

## 3. Materials and Methods

### 3.1. Animal use and ethics

All experiments followed protocols approved by the University of Washington Institutional Animal Care and Use Committees according to the “Guide for the Care and Use of Laboratory Animals” (National Research Council, 2011). Farm porcine hearts were obtained immediately after the animal was euthanized and rinsed in cold oxygenated Tyrode’s buffer.

### 3.2. Protein preparations

Full-length myosin was isolated from animal tissue as previously described.^8^ Briefly, porcine cardiac left ventricular, rabbit psoas, or bovine masseter tissue was minced on ice. Minced tissue was added to extraction buffer (0.3 M KCl, 0.15 M imidazole, 10 mM Na_2_P_2_O_7_, 1 mM MgCl_2_, 2 mM DTT, pH 6.8) and stirred on ice for approximately 30 mins. Excess muscle residue and actin filaments were removed by centrifugation at 260,000 x g for 1 hr. The supernatant was diluted 12X by volume with cold dH_2_O containing 2 mM DTT and left on ice for at least 1.5 hrs to allow myosin to precipitate. Samples were then spun at 44,000 x g to pellet precipitated myosin. Myosin was then dissolved in a high salt solution stopped-flow assay buffer (0.5 M KCl, 20 mM MOPS, 5 mM MgCl_2_, 1 mM DTT, pH 7.0) and was either used for stopped-flow assays as monomeric myosin or converted into synthetic thick filaments. To form synthetic thick filaments, full-length myosin was dialyzed into 0.1 M KCl stopped-flow buffer (20 mM MOPS, 5 mM MgCl_2_, 1 mM DTT, pH 7.0) overnight with at least 2 buffer changes.

### 3.3. Stopped-flow kinetics

Kinetic experiments were performed at 20 °C using a Hi-Tech ‘KinetAsyst’ stopped-flow (SF-61DX2, TgK Scientific, UK). Measurements were made using FRET tryptophan and mant-ATP (Jena Bioscience, NU-202) fluorescence as described before.^1,9^ Briefly, 125 nM myosin was incubated with 2 μM mant-ATP for 1 min before rapidly mixed with 125 μM dark ATP. Transients were collected from 300 – 2500 sec depending on the experimental condition. A retrigger of data collection after the transient was complete for at least 100 s was collected for each condition to ensure baseline stability. Each transient was analyzed individually and the *k*_obs_ values averaged and reported in Tables 1-2 as the mean ± SD of transients as listed.

**Table 1.** Average single-turnover *k*_obs_ values for mavacamten titrations when fit with a single-exponential equation.

| Mavacamten ($\mu\text{M}$) | $k_{\text{obs}}$ ($\text{s}^{-1}$) | |
| --- | --- | --- |
|  | MM | STF |
| 0 | $0.0077 \pm 0.0005$ | $0.0197 \pm 0.0003$ |
| 0.1 | $0.0064 \pm 0.0001$ | $0.0155 \pm 0.0001$ |
| 0.5 | $0.0040 \pm 0.0001$ | $0.0086 \pm 0.0004$ |
| 1 | $0.0028 \pm 0.0001$ | $0.0066 \pm 0.0002$ |
| 10 | $0.0008 \pm 0.0002$ | $0.0014 \pm 0.00003$ |

**Table 2.** Average single-turnover *k*_obs_ values for dATP titrations when fit with a single-exponential equation.

| dATP (%) | $k_{\text{obs}}$ (s<sup>-1</sup>) | |
| --- | --- | --- |
|  | MM | STF |
| 0 | 0.0077 ± 0.0005 | 0.0197 ± 0.0003 |
| 10 | 0.0088 ± 0.0006 | 0.0195 ± 0.0005 |
| 25 | 0.0093 ± 0.00005 | 0.0210 ± 0.0005 |
| 50 | 0.0100 ± 0.00004 | 0.0220 ± 0.0009 |
| 75 | 0.0109 ± 0.00004 | 0.0248 ± 0.0018 |
| 100 | 0.0120 ± 0.00005 | 0.0297 ± 0.00015 |

As discussed in detail in Mohran et al. baseline stability is essential to distinguish a double-exponential fit from a single-exponential with an unstable/drifting baseline.^1^ To this end we use several approaches to minimize drifting baselines and to define any slower process. Firstly, mant. ATP fluorescence was excited via FRET from the protein tryptophan to minimize bleaching of the fluorescence originating from the large excess of mantATP. Secondly, the use of an exciting light reference signal collected just before the observation chamber eliminates any lamp instability and mechanical artifacts from each shot. Finally, after the collection of the transients shown data collection was retriggered on the same sample in the observation chamber to record the base line on each sample.

### 3.4. Data analysis

Data were analyzed using the Kinetic Studio software (TgK Scientific, UK) which provides a least-squares best fit of the with a range of functions including a single-exponential (Fl_t_ = Fl_0_*exp(−*k*_obs_*t) + baseline) or a double-exponential (Fl_t_ = Fl_fast_*exp(−*k*_obs,fast_*t) + Fl_slow_*exp(−*k*_obs,slow_*t) + baseline), as used here. Residuals plots were made by quantifying the difference between the observed signal and the fitted function is provided to evaluate the quality of the fitted function.

## 4. Results

### 4.1. Single-turnover from full-length myosin and synthetic thick filaments exhibit a single population

Full-length monomeric myosin (MM) was prepared to compare mant-ATP single-turnover from skeletal muscle myosin (rabbit psoas) and β-myosin (bovine masseter and porcine left ventricle). MM was dialyzed slowly from 0.5 M KCl to 0.1 M KCl overnight to form synthetic thick filaments (STF). For rabbit psoas and porcine cardiac myosin data from both MM (Figure 1A) and STF (Figure 1B) were well fit with a single-exponential equation with a stable horizontal baseline. For the MM bovine masseter muscle (but not the STF) there was evidence of prolonged slow drift of the baseline. This is assumed to be due to a light scattering artefact caused by slow aggregation of the myosin lasting beyond 1000 s. After correction of the drift the data was well described by a single-exponential (supplemental Fig 1). The analysis yielded rate constants, *k*_obs_, of 0.0051 s^−1^, 0.0071 s^−1^, and 0.0120 s^−1^ respectively for rabbit, bovine and cardiac MM myosins and 0.0159 s^−1^, 0.0178 s^−1^, and 0.0449 s^−1^ respectively for STF.

**Figure 1.**
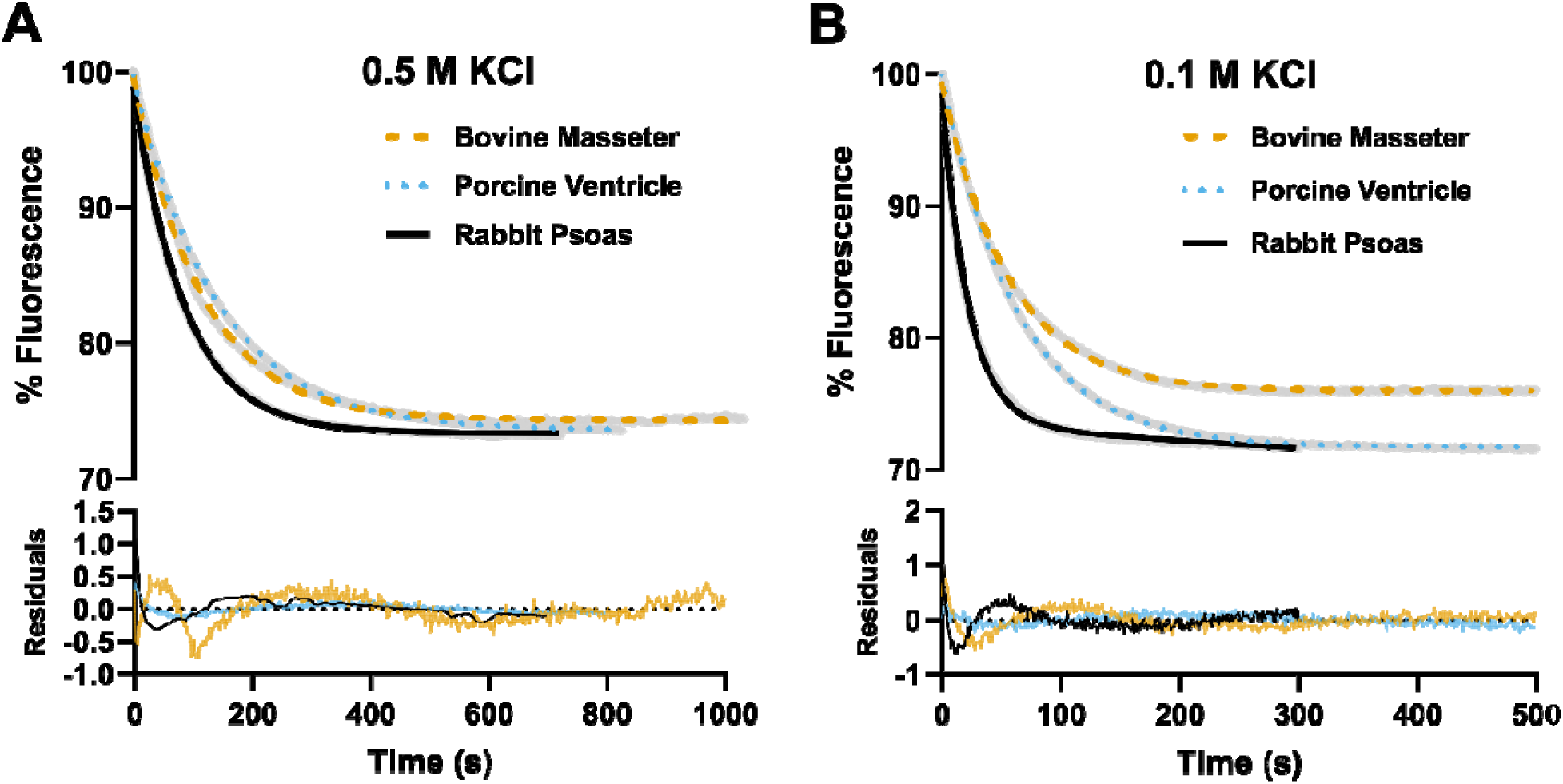
mant-ATP single turnover in (A) full-length myosin at 0.5 M KCl and (B) synthetic thick filaments at 0.1 M KCl from three different tissue sources: bovine masseter, porcine ventricle, and rabbit psoas. Experimental data, gray dots, are fit with a single-exponential equation yielded rate constants, *k*_obs_, of 0.0051 s^−1^, 0.0071 s^−1^, and 0.0120 s^−1^ respectively for full-length myosin; and 0.0159 s^−1^, 0.0178 s^−1^, and 0.0449 s^−1^ respectively for synthetic thick filaments.

Some data does not give a perfect fit to a single-exponential, as revealed by the residual plots, which show some systematic deviation. However, note that the signal change for the reaction is a ~ 25% change of fluorescence while the systematic deviations are ~ 1%. We also analyzed the data using a two-exponential fit, and these are shown for the cardiac MM myosin in Supplemental Figure 2. In the case of the two-exponential fit, the two rate constants gave consistent values but differ by only a factor of two, suggesting the values are unreliable, while the proportion of the two amplitudes (fast:slow) varied widely from 5:1 to 2:3, consistent with parameter redundancy (Supplementary Table 1). In contrast, the single-exponential fits generate consistent values for the rate constant. We therefore do not consider evidence for a second slow phase to be strong unless its amplitude is less than 5% of the major fast phase.

### 4.2. Mavacamten slows the single-turnover rate of full-length myosin and synthetic thick filaments

To determine how an increased IHM population affects single-turnover transients, porcine cardiac MM and STF were treated with a Mavacamten, a well-known myosin inhibitor that stabilizes the IHM and leads to ordering of IHM myosin heads on the thick filament.^10,11^ A range of concentrations from 0 µM (control) to 10 µM (saturating inhibition) were selected to determine the inhibitory constant (K_i_) for both constructs. At all Mavacamten concentrations, transients were best described by a single-exponential decay (Figure 2A,B). At 10 µM, Mavacamten inhibited turnover of mant-ATP from MM by almost 10-fold from 0.0073 s^−1^ ± 0.0001 s^−1^ to 0.0008 s^−1^ ± 0.0002 s^−1^ and STF by almost 15-fold from 0.0197 ± 0.0003 s^−1^ 0.0014 ± 0.00003 s^−1^. Table 1 reports the average values for each condition (n = 3). The Mavacamten titration yielded K_i_ values of 0.48 µM for MM and 0.37 µM for STF.

**Figure 2.**
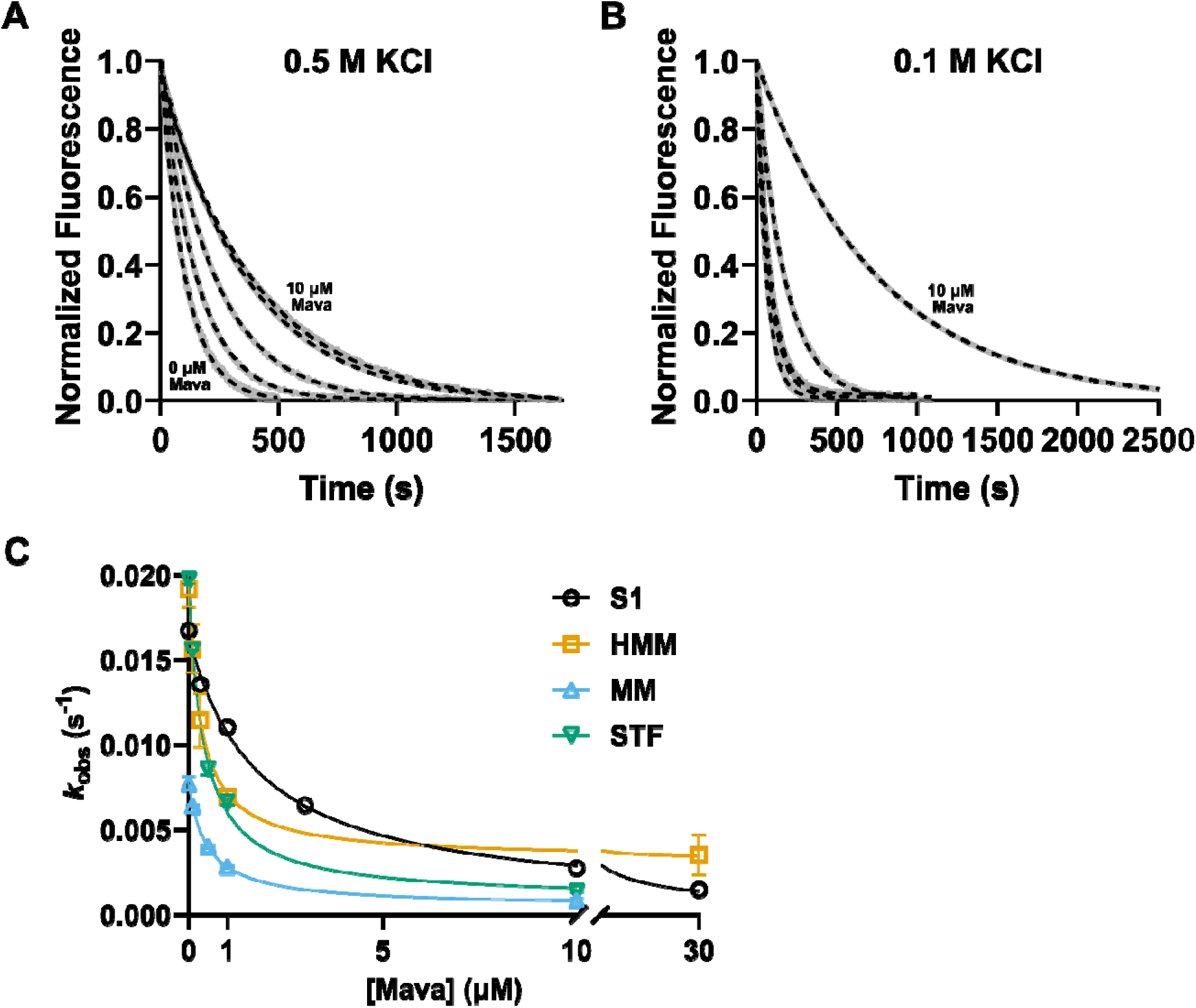
Normalized mant-ATP single-turnover in the presence of mavacamten (Mava) at (A) 0.5 M KCl (monomeric myosin, MM) and (B) 0.1 M KCl (synthetic thick filaments, STF). From left to right: 0 μM, 0.5 μM, 1 μM, 10 μM mavacamten. Experimental data (gray lines) were fit with a single-exponential equation (dashed lines) to obtain *k*_obs_. (C) Plotted *k*_obs_ vs. concentration of mavacamten was fit with a binding isotherm to obtain the inhibitory constant (*K*_i_) of 0.48 μM for MM and 0.37 μM for STF. S1 and HMM data from our previous study (Mohran et al.^1^) is plotted for comparison, *K*_i_ = 1.77 and 0.32 μM respectively. Error bars are ± SD, n = 3-6.

### 4.3. dATP accelerates the single-turnover rate of full-length myosin and synthetic thick filaments

We also replaced some or all of mant-ATP with mant-2’-deoxyATP (mant-dATP) to investigate the effects of the myosin activator on single-turnover transients of MM and STF. dATP has previously been shown to accelerate myosin cycling as well as destabilize the IHM through disrupting ordered thick filament structure.^7,12,13^ Single-turnover transients were collected using a range of dATP concentrations from 0% to 100%, and at all concentrations, transients were best described by a single-exponential decay (Figure 3A, B). dATP increased the single-turnover rate of this single phase in a biphasic manner with a breakpoint at ~50% dATP (Figure 3C), as previously reported for HMM. Table 2 reports average *k*_obs_ values (n = 3). This biphasic behavior was more apparent in STF compared to MM, possibly due to the effects of higher salt concentration disrupting the IHM. Regardless, this may indicate a different behavior of myosins with mant-dATP occupying a single head compared to both, a phenomenon that we previous observed in HMM (the simplest double-headed construct).^1^ Replotting *k*_obs_ vs. the fractional occupancy of two-headed constructs with both heads binding mant-dATP revealed a linear relationship for STF (Figure 3D). However, a non-linear relationship was observed for MM.

**Figure 3.**
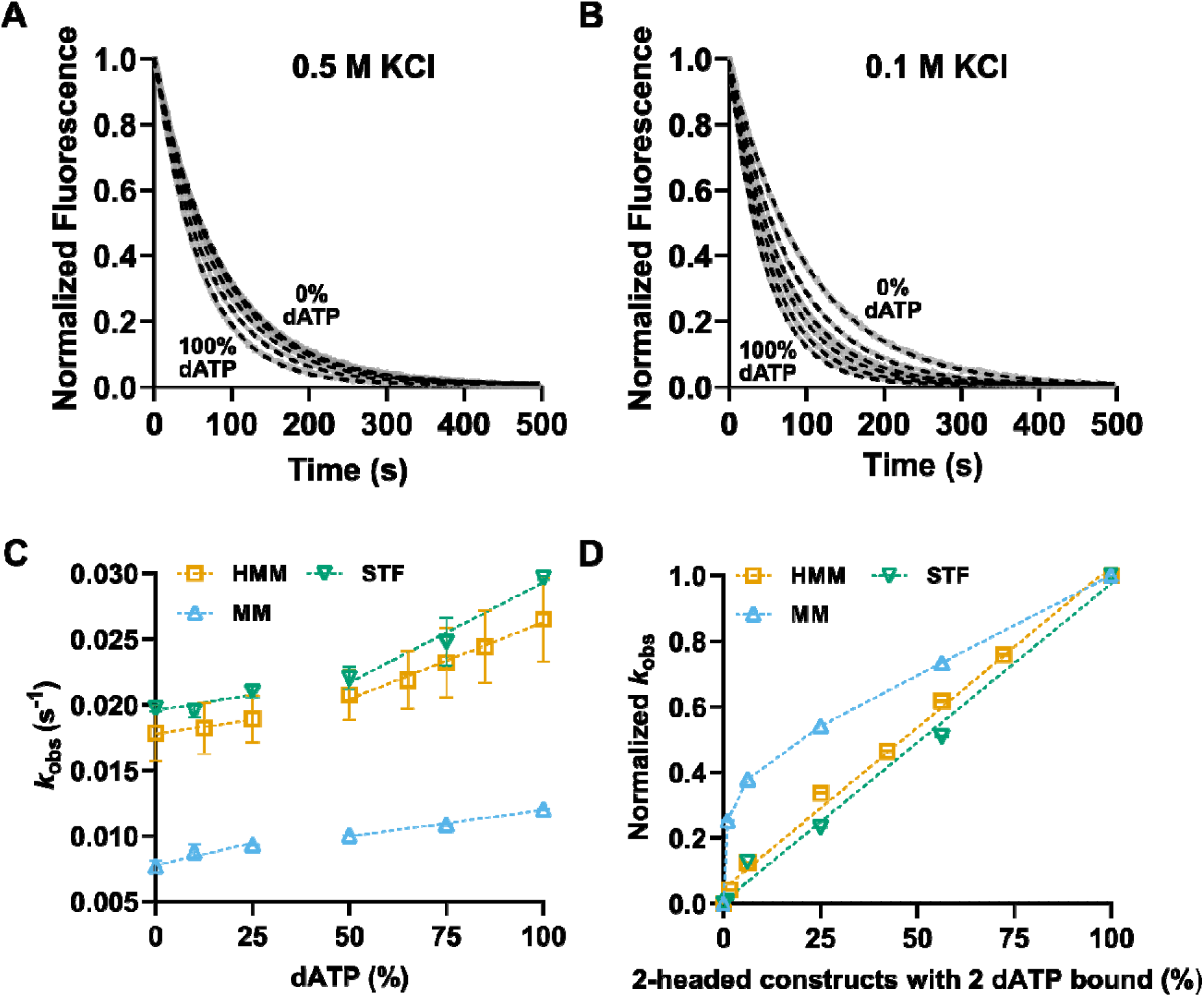
Normalized mant-ATP single-turnover in the presence of dATP at (A) 0.5 M KCl (monomeric myosin) and (B) 0.1 M KCl (synthetic thick filaments). From left to right: 100%, 75%, 50%, 25%, 10%, 0% dATP. Experimental data (gray lines) were fit with a single-exponential equation (dashed lines) to obtain *k*_obs_. (C) Plotted *k*_obs_ vs. % dATP fit with linear lines from 0-50% dATP and 50%-100% dATP for 2-headed constructs. (D) Plotted normalized *k*_obs_ vs. % of 2-headed constructs with 2 dATP bound fit with a linear line from 0-100% dATP for HMM and STF. Data for MM are connected by lines, emphasizing the non-linearity. Error bars are ± SD, n = 3-6.

### 4.4. Porcine cardiac myofibrils exhibit biphasic ATP single-turnover

Isolated myofibrils from porcine left ventricular cardiac tissue were assayed to determine if the presence of whole native filament or sarcomere structure is necessary to produce a single-turnover transient with two populations that have distinct turnover rates. Indeed, these myofibrils produced a fluorescence transient that was best fitted with a double-exponential equation, with rates of *k*_obs,fast_ = 0.0382 ± 0.0043 s^−1^ and *k*_obs,slow_ = 0.0043 ± 0.0010 s^−1^ (Figure 4A,D,E). The relative amplitudes of the fast and the slow phase (A_fast_:A_slow_) was 76:24 (Figure 4B,C). When the myofibrils were pre-incubated with 0.5 µM Mavacamten, the relative amplitude was shifted towards the slow phase (56:44, Figure 4B,C). Additionally, the rate of the fast phase was significantly decreased (*k*_obs,fast_ = 0.0263 ± 0.0016 s^−1^, *p* < 0.05, Figure 4D). Replacing mant-ATP with mant-dATP produced fluorescence transients that were best fit with a single-exponential where the slow phase observed in the no drug (ND) and Mavacamten conditions was completely abolished (Figure 4A). Furthermore, the rate of the fast phase was slightly elevated (*k*_obs,fast_ = 0.0455 ± 0.0026 s^−1^, *p* = 0.07, Figure 4D) as seen for isolated myosin (Figure 3).

**Figure 4.**
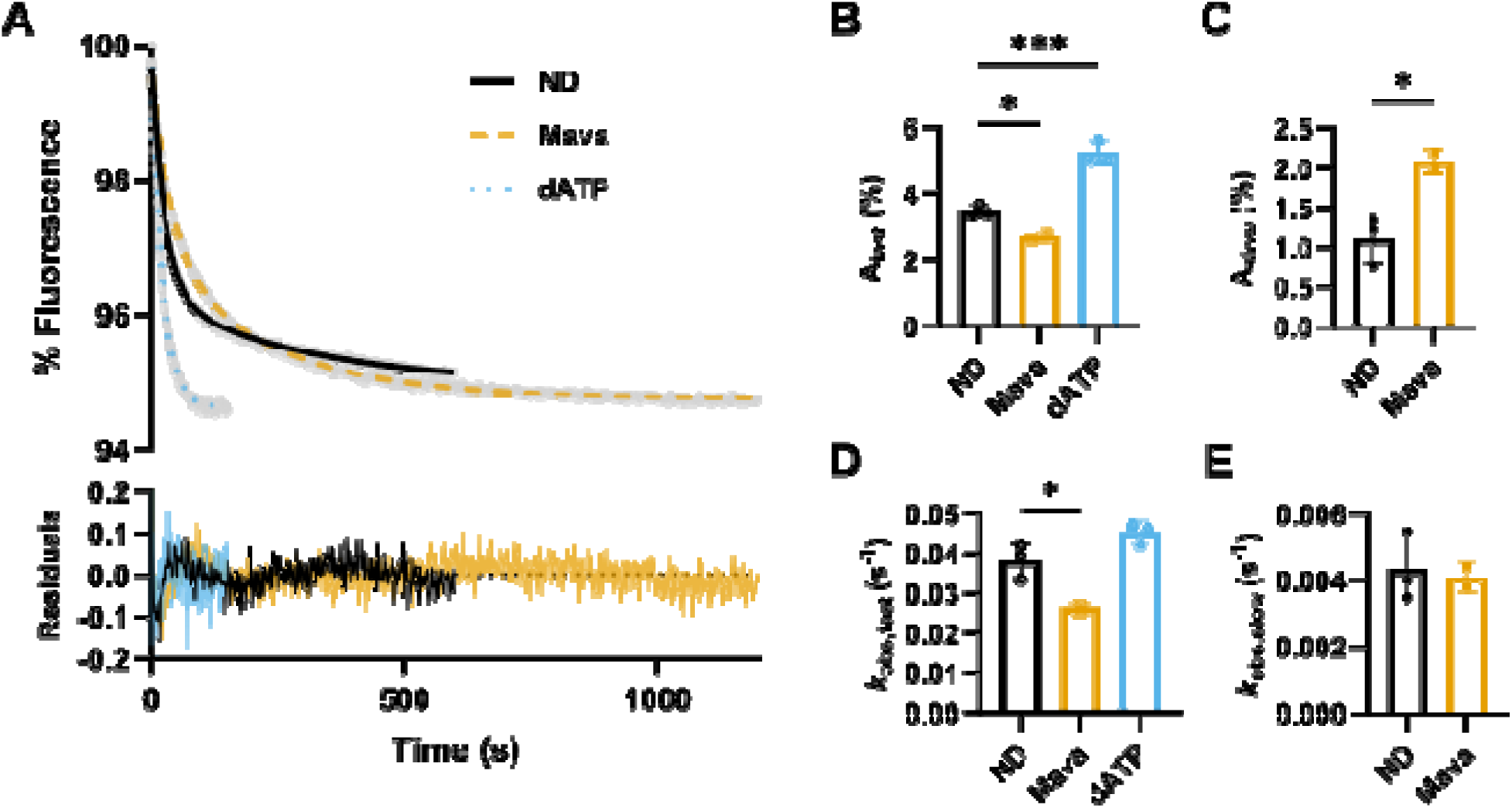
(A) mant-ATP single-turnover in pig cardiac myofibrils. Experimental data (gray lines were fit with a double-exponential equation in the no drug (ND) and 0.5 μM mavacamten (Mava) conditions and a single-exponential equation in the 100% dATP condition (dATP). (B-E) Fit parameters of the fast and slow phases for each condition. Error bars are ± SD, n = 2-3. * *p* < 0.05, *** *p* < 0.001, student’s t-test.

## 5. Discussion

In this work we have extended our earlier studies on native HMM to full-length striated muscle myosin and myofibrils. For myosin, as for HMM, we find no evidence of a slow phase in the mant-ATP displacement assay which has been widely used to define the SRX state of myosin. Significantly, this is true for both monomeric myosin at 0.5 M KCl (where the high salt concentration is expected to destabilize the SRX state), and at 0.1 M KCl where myosin typically forms synthetic bipolar filaments much shorter than native thick filaments. This result is in direct contrast to the study of Gollapudi *et al.* who did report a significant slow phase with synthetic myosin filaments.^14^ but we did point out in our earlier study that the SRX assay of Gollapudi *et al.* was not consistent with their own ATPase data reported at the same time.^1^ The Gollapudi *et al.* observations are consistent with a large body of data using short HMM expressed in C2C12 cells. These consistently report fast and slow phases in the displacement reaction. The short HMM consists of S1 ligated with variable lengths of the S2 tail (2 – 25 heptads) followed by a leucine zipper to ensure a stable dimer. The short HMM constructs used contrasts with the 42-heptad construct used by Cail *et al.* and the 42-heptad native protein in Mohran *et al*.^1^ It is therefore possible that the differences in the results observed by the different groups could be due, to some degree, to the different HMM preparations used.

Recent reports from Goluguri *et al*. and Duno-Miranda *et al*. report a two phase mant-ATP displacement reaction.^15,16^ We currently have no explanation for the difference between the two studies showing one or two phases. In contrast, our myosin and HMM data is consistent with theoretical consideration in which a slow phase cannot be present if the SRX state is in equilibrium with the DRX state on the time scale of the assay, e.g. ~15 minutes. This puts a severe limit on any model of the relationship between the SRX and DRX states. The two states must be distinct populations of myosin heads that do not interconvert over the 15-minute time scale of the assay.

In addition to the basic observations, we report that Mavacamten inhibits the mant-ATP turnover rate constant for both monomeric and STF with *K*_i_’s (0.55 & 0.38 µM respectively). This is similar to that we reported for HMM (0.32 µM) but much tighter than the Ki for single headed S1 (1.77 µM). The *K*_i_’s for S1, HMM and STF were all measured at the same KCl concentration and can be compared. The five-fold tighter affinity of mavacamten for HMM and STF than for S1 suggest there is some cooperativity between the two heads in the binding of Mavacamten, although the binding isotherms are too tight to reveal any sigmoidicity.

A similar interaction between the two myosin heads was apparent for dATP binding to HMM in our previous study.^1^ A plot of k_obs_ vs % mant-dATP present showed an acceleration of the rate constant of displacement but with two distinct slopes, one at < 50 % dATP and a greater one for dATP > 50%. If the probability of a myosin head having a dATP bound is α then the probability of both heads having dATP bound is α^2^. Replotting the data as the fraction of HMM expected to have 2 dATP bound gave a single slope which is consistent with the NTPase acceleration requiring both heads to have dATP bound. A similar behavior is reported here for STF (Figure 3). S1 showed no such effect, as was also true of monomeric myosin at high KCl. This again provides evidence for an interaction between the two heads of myosin in either STF or HMM preparations but not in S1 or MM at high salt. The presence of dATP vs ATP accelerates the *k*_obs_ of the displacement reaction, but the effect is much stronger when both heads bind dATP.

In summary we see no direct evidence of the SRX state as classically defined for HMM STF or for monomeric myosin. Also, Mavacamten inhibits, while dATP accelerates the turnover of ATP by all three constructs and for S1. The difference between S1 versus HMM and STF is that there is clear evidence for cooperativity between the two heads of HMM and STF when binding either mavacamten or dATP. This is consistent with the structurally defined IHM. At high salt concentrations monomeric myosin is intermediate between S1 and HMM. Monomeric myosin binds Mavacamten with a *K*_i_ three times tighter than that of S1 but almost 2-fold weaker than HMM or STF. This may simply be the effect of ionic strength on the binding of mavacamten or dATP, but an effect on head-head interactions cannot be eliminated since ionic strength is expected to weaken the interactions in IHM. In contrast, dATP shows little evidence of head-head interaction in its acceleration of the turnover.

The results with myofibrils are quite different to those with HMM, STF and myosin. Here we observed a consistent slow phase, which is similar to that reported earlier by Walklate for fast skeletal, slow skeletal and cardiac myofibrils.^7^ The myofibrils are intrinsically more variable than the purified proteins and the total amplitude is observed is much smaller (~4-5% compared to 20-25% for isolated myosin), making the detailed analysis more problematic, The slow phase represents only 1% of the total signal with a rate constant that is 10% of the fast phase. The fast phase is larger and thus easier to define. The fast phase rate constant is reduced by 5 µM Mavacamten, but only by ~ 30%, which is much less than that seen for the proteins. 100% dATP accelerates the fast phase rate constant by ~ 10%, similar to that seen in the proteins. It is also clear that 100% dATP eliminates the slow phase completely. It is the contrast between the +/- dATP transient that makes it clear that the slow phase in the absence of dATP is genuine. Detailed titrations of either Mavacamten or dATP with myofibrils, as was done for HMM and STF, are challenging as the slow phase amplitude is small, so evaluating change in the slow phase rate constant or amplitude proved futile.

So, what is the source of the slow phase in the myofibril assay? As stated above, a slow phase of this type cannot be generated by two forms of myosin in equilibrium with each other on the minute time scale, they must represent distinct populations of myosins which are not interconverting. There are several possible sources of distinct populations in the sarcomere. It is known that the D, C (and P) zones are distinct because of the presence of MyBP-C in the C-zone. Furthermore cryo-EM images of the relaxed C-zone indicate three populations of myosin crowns within the 43 nm repeat of the thick filament, with each showing different degrees of order/disorder. Thus, there are several distinct populations of myosin heads having distinct locations within the thick filament and they therefore do not form single population. The myosin in each population can have its own properties that affect, among other things, the ATPase rate and a distinct partitioning between the “On” and “Off” states.

For a single population of myosin heads in equilibrium between On and Off states, then

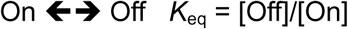

If On heads turn over ATP at *k*_On_ = 0.05 s^−1^ and the Off heads turn over ATP at *k*_Off_ = 0 s^−1^, then the net displacement rate constant (*k*_displacement_) is the fraction of the total heads in the On-state (α) multiplied by *k*_On_.

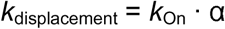

If *k*_fast_ = 0.04 s-1 and *k*_slow_ −0.004 s-1 then the fast population has α_fast_ = *k*_fast_/*k*_On_ = 0.04/0.05 = 0.8 or 80% On. For the slow phase α_slow_ = *k*_slow_/*k*_On_ = 0.004/0.05 = 0.08 or 8% On. In this model the relative amplitude of the two phases defines the size of the two populations while each *k*_obs_ represents the intrinsic turnover rate of a myosin head times the fraction of the population available (i.e., not in an IHM).

It is tempting to assign these two populations to the D and C zones of the thick filament but there is no direct evidence for such a distinction between the two phases. Possible indirect evidence is that in cryo-EM and cryo-tomography the C-zone of the thick filament shows some degree of order in the myosin heads while, in contrast, little order is apparent in the D zone. The single fluorescent ATP molecule displacement assay would argue against such a simple assignment as these indicate two populations of myosin turning over ATP at different rates in both the C and D zones, with differing proportions of fast and slow components.^17,18^ Indeed our numbers for control, Mavacamten and dATP in myofibril preparations for the mant-NTP displacement assay are in broad agreement with those reported in Pilagov *et al.* using the single fluorescent ATP molecule displacement assay where fast and slow rates were observed in all three thick filament zones. Thus, it cannot be assumed that there are only two populations of myosins, just that the analysis can reveal the presence of at least two populations.

In conclusion, we find no evidence of the SRX state as originally defined by a slow displacement of mant-ATP from S1, or HMM or myosin in either monomeric form or in synthetic thick filaments. This does not mean that an IHM does not exist in our preparations, just that any IHM is in rapid equilibrium with free, DRX heads. A careful evaluation of turnover rates for S1 vs two-headed constructs can provide evidence of the presence of IHM, as recently explored by Cail et al. Our data do provide evidence of interaction between the two heads of myosin from the Ki of Mavacamten binding and the greater effect of dATP when both heads have dATP bound. Finally relaxed myofibrils do show the presence of two or more populations of myosins in the thick filament sarcomere which are not seen for isolated myosin. Additional components of the thick filament or sarcomere are required to generate distinct populations of myosins with different fractions of the heads available to turn over ATP.

## Author contributions

Designed research, MAG and MR

Performed experiments, KYK, MAG, KZR, AMA, TWW, and TSM

Analyzed data, KYK and MAG

Manuscript writing, KYK, MAG, and MR

Resources and funding, MR, MAG, and JDP

All authors contributed intellectually to discussions of the results and reviewed and edited the paper.

## Data availability

Data is available upon request through.

