## Supplemental Figures for "Revisiting the SRX paradox for myosin and myofibrils"

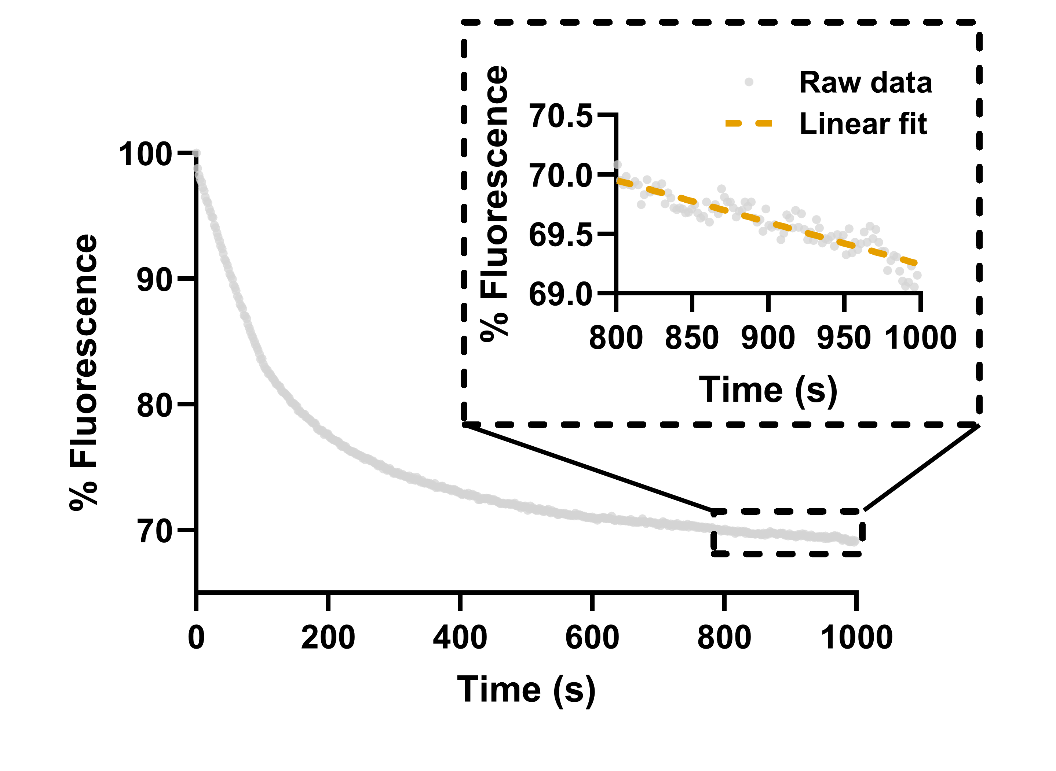

**Supplemental Figure 1.** Mant-ATP single turnover of monomeric myosin (0.5 M KCl) from bovine masseter tissue. **(Inset)** Raw data (gray dots) show a linear baseline drift of -0.00076 s^-1^ which is corrected in Figure 1A.

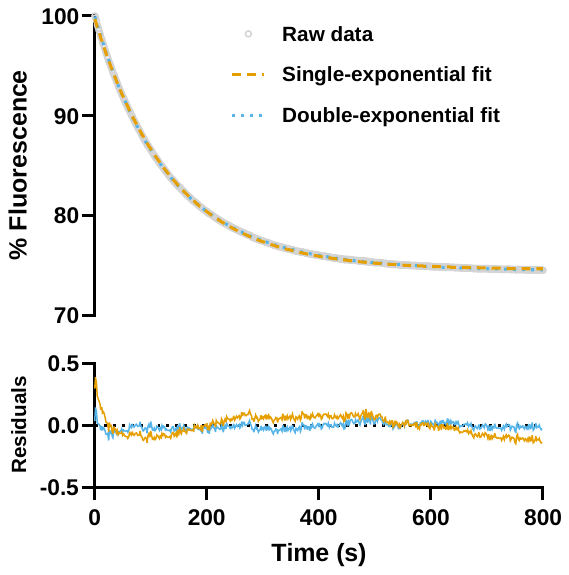

**Supplemental Figure 2.** Single- and double-exponential fits of porcine cardiac myosin 0.5 M KCl (monomeric myosin) no drug (ND) replicate 1. Single-exponential fit with parameters A = 25.67% and *k*_obs_ = 0.00734 s^-1^. Double-exponential fit with parameters A_fast_ = 18.02%, A_slow_ = 8.30%, *k*_obs,fast_ = 0.00883 s^-1^, and *k*_obs_ = 0.00445 s^-1^. Relative *k*_obs_ parameters of the double-exponential fit are less than 3-fold different, which may lead to parameter redundancy.

**Supplemental Table 1.** Single- and double-exponential fits for traces at 0.5 M KCl (monomeric myosin).

| **Condition** | **Replicate** | **Single-exponential** | | **Double-exponential** | | | |
| --- | --- | --- | --- | --- | --- | --- | --- |
|  |  | **A** | ***k*_obs_** | **A_fast_** | **A_slow_** | ***k*_obs,fast_** | ***k*_obs,slow_** |
| **ND** | 1 | 25.67 | 0.00734 | 18.02 | 8.3 | 0.00883 | 0.00445 |
|  | 2 | 24.55 | 0.00772 | 11.23 | 14.33 | 0.00997 | 0.00588 |
|  | 3 | 26.15 | 0.00726 | 11.38 | 15.49 | 0.01004 | 0.00572 |
|  | 4 | 26.16 | 0.00734 | 11.23 | 14.33 | 0.00997 | 0.00588 |
|  | 5 | 40.58 | 0.00809 | no fit |  |  |  |
|  | 6 | 50.35 | 0.00743 | 30.24 | 21.91 | 0.00993 | 0.00476 |
|  | 7 | 49.75 | 0.00756 | 42.73 | 8.73 | 0.00857 | 0.0032 |
| **10% dATP** | 1 | 23.99 | 0.0081 | 0.897 | 23.47 | 0.03796 | 0.00792 |
|  | 2 | 24.79 | 0.00917 | 5.62 | 19.14 | 0.00917 | 0.00917 |
|  | 3 | 23.6 | 0.00911 | 5.07 | 18.51 | 0.00911 | 0.00911 |
| **25% dATP** | 1 | 31.11 | 0.00998 | 3.24 | 29.56 | 0.06199 | 0.00939 |
|  | 2 | 29.35 | 0.0093 | 29.98 | -2.18 | 0.00899 | 0.0008 |
|  | 3 | 27.6 | 0.00934 | -13.43 | 40.79 | 0.01284 | 0.01031 |
| **50% dATP** | 1 | 28.93 | 0.01008 | -10.64 | 39.44 | 0.01316 | 0.01081 |
|  | 2 | 29.21 | 0.01004 | 0.43 | 29.14 | 0.23412 | 0.01 |
|  | 3 | 28.97 | 0.01001 | -5.82 | 34.74 | 0.01233 | 0.01036 |
| **75% dATP** | 1 | 29.87 | 0.01087 | -3.36 | 33.2 | 0.01301 | 0.01107 |
|  | 2 | 29.81 | 0.01092 | -4.72 | 34.46 | 0.01398 | 0.01129 |
|  | 3 | 35.74 | 0.01084 | -3.15 | 38.87 | 0.01238 | 0.01096 |
| **100% dATP** | 1 | 32.57 | 0.01197 | -10.84 | 43.27 | 0.0154 | 0.01273 |
|  | 2 | 31.23 | 0.01206 | 32.42 | -3.42 | 0.01147 | 0.00132 |
|  | 3 | 30.84 | 0.01205 | -13.84 | 44.31 | 0.01771 | 0.01351 |
| **0.1 μM mava** | 1 | 28.69 | 0.00643 | 23.12 | 6.29 | 0.00769 | 0.00348 |
|  | 2 | 27.4 | 0.00629 | 13.52 | 14.32 | 0.00831 | 0.00498 |
|  | 3 | 27.1 | 0.00647 | 0.84 | 26.52 | 0.02291 | 0.00635 |
| **0.5 μM mava** | 1 | 25.05 | 0.00397 | 0.52 | 24.96 | 0.09682 | 0.00395 |
|  | 2 | 24.02 | 0.00396 | 0.39 | 23.96 | 0.13422 | 0.00395 |
|  | 3 | 26.27 | 0.00416 | 5.6 | 20.88 | 0.0059 | 0.00383 |
| **1 μM mava** | 1 | 23.57 | 0.00292 | 0.47 | 23.51 | 0.08673 | 0.00291 |
|  | 2 | 22.94 | 0.0028 | 0.97 | 22.77 | 0.06434 | 0.00277 |
|  | 3 | 22.03 | 0.00274 | 0.36 | 21.99 | 0.11667 | 0.00273 |
| **10 μM mava** | 1 | 27.82 | 0.001 | 6.79 | 23.48 | 0.00263 | 0.00073 |
|  | 2 | 19.74 | 0.00068 | 3.74 | 16.4 | 0.00109 | 0.0006 |
|  | 3 | 23.43 | 0.00079 | 13.11 | 12.64 | 0.00116 | 0.0004 |

**Supplemental Table 2.** Single- and double-exponential fits for traces at 0.1 M KCl (synthetic thick filaments).

| **Condition** | **Replicate** | **Single-exponential** | | **Double-exponential** | | | |
| --- | --- | --- | --- | --- | --- | --- | --- |
|  |  | **A** | ***k*_obs_** | **A_fast_** | **A_slow_** | ***k*_obs,fast_** | ***k*_obs,slow_** |
| **ND** | 1 | 38.07 | 0.01908 | 7.48 | 30.2 | 0.01982 | 0.01982 |
|  | 2 | 36.12 | 0.01996 | 9.54 | 26.58 | 0.02 | 0.01995 |
|  | 3 | 38.81 | 0.01868 | 7.56 | 30.84 | 0.01941 | 0.01941 |
| **10% dATP** | 1 | 36.48 | 0.01901 | -19.35 | 55.66 | 0.02389 | 0.0205 |
|  | 2 | 36.47 | 0.01984 | 0.36 | 36.31 | 0.12657 | 0.01952 |
|  | 3 | 42.33 | 0.01976 | no fit |  |  |  |
| **25% dATP** | 1 | 31.79 | 0.0208 | no fit |  |  |  |
|  | 2 | 34.42 | 0.02061 | 0.59 | 34.09 | 0.10575 | 0.0204 |
|  | 3 | 32.03 | 0.02153 | 0.24 | 31.94 | 0.19355 | 0.02146 |
| **50% dATP** | 1 | 34.7 | 0.02105 | no fit |  |  |  |
|  | 2 | 34.62 | 0.02196 | 2.61 | 31.78 | 0.02246 | 0.02245 |
|  | 3 | 39.88 | 0.02263 | 39.94 | -1.32 | 0.0225 | 0.00028 |
| **75% dATP** | 1 | 33.08 | 0.02271 | 33.15 | -0.51 | 0.02261 | 0.00081 |
|  | 2 | 35.46 | 0.02549 | 1.91 | 33.55 | 0.02549 | 0.02549 |
|  | 3 | 33.09 | 0.02617 | 1.75 | 31.34 | 0.02617 | 0.02617 |
| **100% dATP** | 1 | 35.56 | 0.0295 | 2.2 | 33.56 | 0.0295 | 0.0295 |
|  | 2 | 34.05 | 0.02967 | 2.03 | 32.02 | 0.02968 | 0.0967 |
|  | 3 | 34.3 | 0.0298 | 1.55 | 32.75 | 0.0298 | 0.0298 |
| **0.1 μM mava** | 1 | 35.92 | 0.01563 | 2 | 33.92 | 0.01563 | 0.01563 |
|  | 2 | 34.4 | 0.01561 | 3.83 | 30.57 | 0.01566 | 0.01561 |
|  | 3 | 33.76 | 0.01553 | 3.16 | 30.59 | 0.01553 | 0.01553 |
| **0.5 μM mava** | 1 | 33.17 | 0.009 | 1.1 | 32.97 | 0.19282 | 0.00892 |
|  | 2 | 32.95 | 0.00849 | -11.94 | 44.79 | 0.01044 | 0.00895 |
|  | 3 | 30.04 | 0.0084 | -19.69 | 49.53 | 0.01084 | 0.00925 |
| **1 μM mava** | 1 | 27.02 | 0.00709 | 19.5 | 7.52 | 0.00709 | 0.00709 |
|  | 2 | 23.27 | 0.00667 | 24.24 | -5.06 | 0.00615 | 0.00034 |
|  | 3 | 21.14 | 0.00667 | -44.86 | 65.67 | 0.00891 | 0.00807 |
| **10 μM mava** | 1 | 19.91 | 0.00144 | 1.44 | 20.02 | 0.08897 | 0.00139 |
|  | 2 | 18.65 | 0.00137 | 19.27 | -1.27 | 0.00134 | 0.00027 |
|  | 3 | 17.08 | 0.00134 | 18.09 | -1.35 | 0.00129 | 0.00049 |
